# Germline Sex Determination regulates sex-specific signaling between germline stem cells and their niche

**DOI:** 10.1101/2021.04.30.442141

**Authors:** Pradeep Kumar Bhaskar, Sheryl Southard, Kelly Baxter, Mark Van Doren

## Abstract

The establishment of sexual identity in germ cells is critical for the development of male and female germline stem cells (GSCs) and production of sperm vs. eggs. Thus, this process is essential for sexual reproduction and human fertility. Germ cells depend on signals from the somatic gonad to determine their sex, but in organisms such as flies, mice and humans, the sex chromosome genotype of the germ cells is also important for germline sexual development. How somatic signals and germ cell-intrinsic cues act together to regulate germline sex determination is a key question about which little is known. We have found that JAK/STAT signaling in the GSC niche promotes male identity in germ cells and GSCs, in part by activating expression of the epigenetic reader Phf7. We have also found that JAK/STAT signaling is blocked in XX (female) germ cells through the intrinsic action of the sex determination gene *Sex lethal*, which preserves female identity. Thus, an important function of germline sexual identity is to control how GSCs respond to signals in their niche environment.

## Introduction

Sexual dimorphism, the differences between the sexes, manifests as distinct developmental programs in males and females, resulting in sex-specific anatomy, physiology and behavior. Nowhere is this more important than in the germline, which is responsible for producing the sex-specific gametes necessary for perpetuation of the species. However, the process of sex determination in the germline is still poorly understood in most animals. One key aspect of sex determination is whether it is established autonomously by the germ cell’s own sex chromosome constitution, non-autonomously via signals from somatic cells, or both. In some animals, the sex of the soma is sufficient to control the sex of the germline, as germ cells are able to follow the correct developmental path (spermatogenesis or oogenesis) regardless of the sex of the soma (Hilfiker-Kleiner et al., 1994; Blackler, 1965; Yoshizaki et al., 2011). However, in organisms such as Drosophila and humans, germ cell development fails if the “sex” of the germline does not match the “sex” of the soma. For example, in Drosophila, XX (normally female) germ cells transplanted into an XY (male) somatic environment are lost during development (Van Deusen, 1977), while germ cells in sex-transformed “XX males” are atrophic and fail to develop into sperm (Sturtevant, 1945; Nöthiger et al., 1989) (Figure 2B). Similarly, XY germ cells present in a female somatic environment do not enter oogenesis but instead form germline tumors (Schüpbach, 1982, 1985; Steinmann-Zwicky et al., 1989). In humans, Turner’s Syndrome patients have an X0 genotype, and the soma follows a female developmental program due to the lack of a Y chromosome. However, these patients are infertile due to germline loss and incomplete oogenesis (Ye et al., 2020). Thus, the presence of a single X chromosome in the germline is thought to be incompatible with female germline development. Similarly, Klinefelter’s patients (XXY) develop somatically as males but are also infertile, and rare patches of spermatogenesis observed in testes of these individuals are due to germ cells having lost one X chromosome (now XY) (Deebel et al., 2020). Thus, in both flies and humans, the sex chromosome genotype of the germ cells has a strong effect on proper germline sexual development, indicating that germline autonomous cues combine with non-autonomous cues from the soma to regulate this process.

**Figure 1:**
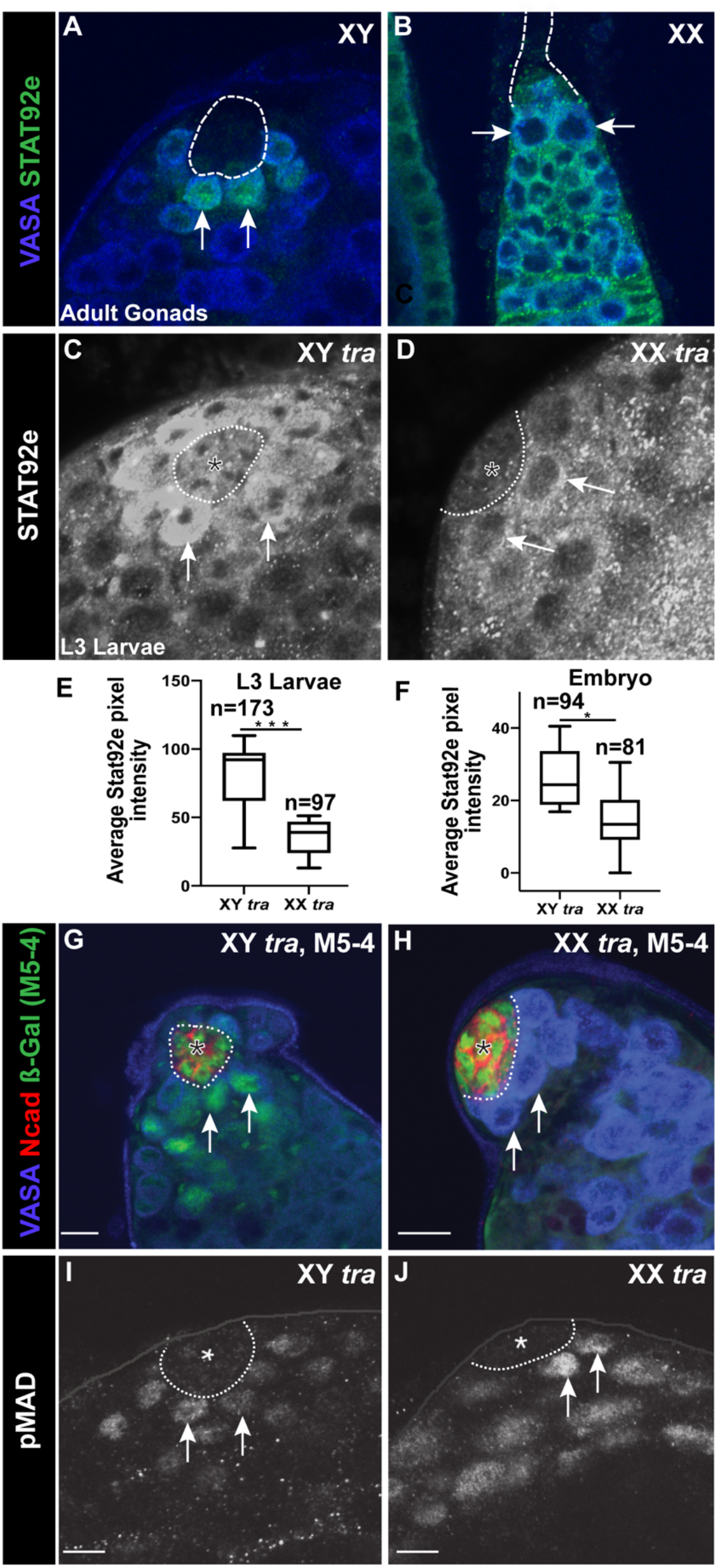
XX GSCs show a decreased response to JAK-STAT signaling from the niche. Antibody staining as indicated in figure. Anti-Vasa labels germ cells. GSC niches outlined with dashed line. (A-B) Anti-Stat92E (green) labeling in adult gonads. Note that GSCs (e.g., arrows) exhibit strong immunoreactivity in males (A) but not in females (B). (C, D) Anti-Stat92E immunostaining of XY *tra* or XX *tra* L3 testes. Note the reduced immunostaining in XX *tra* testes (D). (E, F) Box and whiskers plot of the mean Stat92E pixel intensity per testis for XY *tra* or XX *tra* L3 testes (E) or stage 15-16 embryos (F). (G, H) M5-4 enhancer trap in XY *tra* and XX *tra* testes. Note the loss of M5-4 staining in germ cells of XX testes (H). (I, J) pMad expression in XY *tra* L3 testes (I), XX *tra*, L3 testes (J).

**Figure 2:**
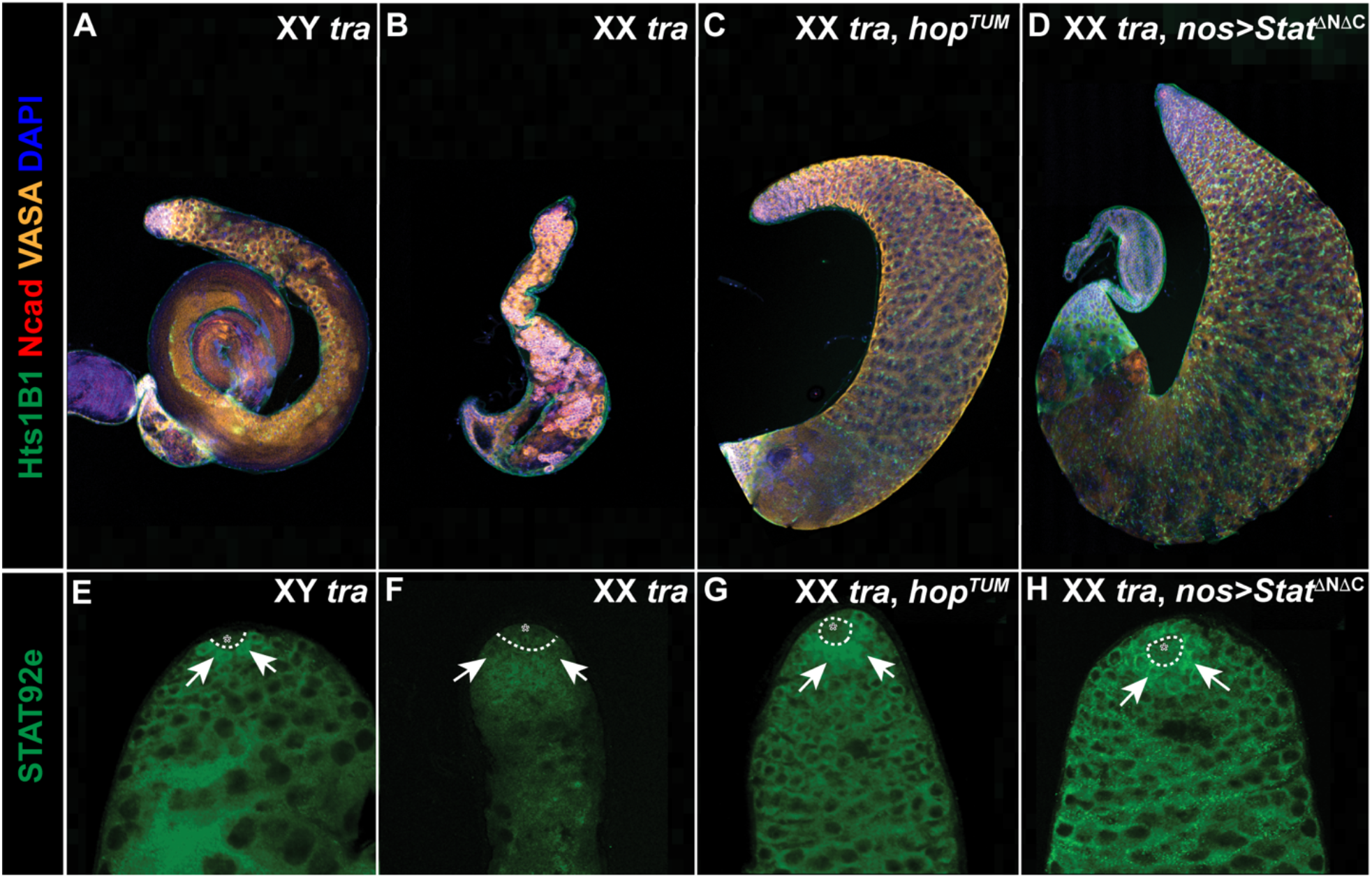
Restoring JAK-STAT signaling in XX *tra* testes partially rescues germline depletion. Antibody staining as indicated. (A-D) Anti-Vasa labels germ cells, Anti-Hts1B1 labels the fusome, Anti-NCad labels the hub. (A) XY *tra* (control) testis, B) XX *tra* testes, (C) XX *tra* testes expressing activated JAK, *hop^TUM^*, and (D) XX *tra* testes expressing activated STAT92E in germ cells, nos>Stat92E^ΔNΔC^. (E-H) Anti-Stat92E labeling (Green) of the same genotypes as indicated in (A-D). Hubs are indicated by dashed lines and representative GSCs indicated by arrows. Note that the decreased anti-Stat92E immunostaining observed in (F) is rescued by activation of the JAK/STAT pathway in (C) and (D).

In Drosophila, *Sex lethal (Sxl)* is the key gene acting autonomously in the germline to promote female sexual identity. *Sxl* encodes an RNA binding protein that acts in regulating both alternative mRNA splicing and translational control (Bashaw and Baker, 1997; Bell et al., 1988; Keyes et al., 1992; Kelley et al., 1997; Gebauer et al., 1998). Loss of *Sxl* specifically from germ cells disrupts oogenesis and causes germ cells to produce ovarian tumors, similar to XY germ cells developing in a female soma (Schüpbach, 1985). Further, expression of *Sxl* in XY germ cells is sufficient to allow them to complete oogenesis when present in a female soma (Hashiyama et al., 2011). Interestingly, *Sxl* is also the key “switch” gene that regulates female identity in the soma (Cline, 1979). In both the soma and the germline, *Sxl* expression depends on the presence of two X chromosomes, but the genetics of *Sxl* activation in the female soma are different from that in the germline (Granadino et al., 1993; Steinmann-Zwicky, 1993). Further, the targets for regulation by Sxl in the soma, *transformer (tra)*, which regulates female somatic identity, and *male specific lethal 2*, which regulates X chromosome dosage compensation in the soma, are not required in the germline (Bachiller and Sanchez, 1986; Marsh and Wieschaus, 1978). Thus, the way in which Sxl acts to control female sexual identity in the germline is largely unknown, although one target for Sxl regulation in the germline has been identified (Primus et al., 2019).

How signals from somatic cells also act to control germline sexual identity is similarly unknown. Previously, we have found that the JAK/STAT (Janus kinase/ signal transducer and activator of transcription) pathway acts as an important signal from the soma to promote male identity in the germline (Wawersik et al., 2005). Initially, in the embryonic gonad, male somatic cells express JAK/STAT ligand(s) which promote male gene expression and behavior in the germline (Wawersik et al., 2005; Sheng et al., 2009). However, the JAK/STAT pathway is eventually activated in the gonads of females as well, where it is important for essential somatic cell types (the escort cells) (Decotto and Spradling, 2005). Why the presence of the JAK/STAT pathway in the ovary does not masculinize the germ cells has remained unknown.

One place where germline sexual dimorphism is particularly apparent is in the germline stem cells (GSCs). GSCs are responsible for the continuous production of germ cells that enter gametogenesis and produce the large numbers of gametes necessary for full fertility. In many animals, GSCs are present in both males and females, yet their behavior, and the processes of spermatogenesis vs. oogenesis, exhibit clear sexual dimorphism (Casper and Van Doren, 2006; Spradling et al., 2011). Further, mammals like mice and humans only have germline stem cells in the testis, while the ovary contains a pre-determined set of developing oocytes. Thus, the differences between the sexes are even more extreme in these animals. How germline sex determination leads to such dramatic differences in GSC behavior and potential is largely unexplored. Interestingly, the JAK/STAT pathway is also a key regulator of GSC behavior in Drosophila males but is not required in female GSCs (Kiger et al., 2001; Tulina and Matunis, 2001; Leatherman and Dinardo, 2010; Decotto and Spradling, 2005; Chen et al., 2018). Thus, the JAK/STAT pathway may provide a link between germline sex determination and GSC identity and behavior.

Here we show that one way in which *Sxl* acts to promote female identity in the germline is by blocking reception of the JAK/STAT signal in male GSCs. Further, we show that the epigenetic reader *Plant homeodomain-containing factor 7 (Phf7)*, an important regulator of male identity in the germline (Yang et al., 2012), is a direct downstream target of JAK/STAT signaling in male germ cells. Thus, a key aspect of how germline autonomous information interacts with non-autonomous signals from the soma is by influencing how GSCs interact with their niche environment.

## Results

### XX germ cells exhibit decreased JAK/STAT signaling

To explore how the somatic environment and germline autonomous cues combine to control germline sexual development, we first explored the development of XX germ cells in a male somatic environment. *tra* is essential for female identity in the soma, and *tra* mutant animals exhibit a robust female to male transformation, but *tra* has no known role in males and XY *tra* animals are indistinguishable from wt males (Sturtevant, 1945; Watanabe and Onishi, 1975). However, since *tra* is not required in the female germline (Marsh and Wieschaus, 1978), XX *tra*-mutants allow us to study otherwise normal XX germ cells in a male somatic environment (hereafter referred to as XX males). Previously, it has been shown that the germline in XX males is severely atrophic, with depleted germline and a lack of differentiating germ cells (Steinmann-Zwicky et al., 1989) (Figures S1A, S1B). We examined gonad development in XX males and found that soma-germline interaction and formation of the male germline stem cell niche appeared completely normal until the 3^rd^ instar larval (L3) stage (Murray (nee Southard) S, 2011). However, L3 gonads of XX males exhibited an atrophic germline (Compare Figures S1D to S1C) with a reduced number of GSCs (avg. # GSCs: control male = 10.9 + 1.75, n=16; XX male = 6.85 + 1.5, n=13). One of the main differences between XY and XX animals is that control XX females and XX males express the sex determination factor Sxl in the germline, while XY males do not (Figures S1E–S1G). In addition, XY individuals expressing ectopic TraF protein (XY females) (Figure S1H) also fail to express Sxl, indicating that expression of Sxl in the germline is dependent only on the germ cell genotype and is independent of somatic cues.

Another important sex-specific quality of the GSCs is their requirement for different signals from the surrounding niche. Male GSCs require signaling through the JAK/STAT pathway for their proper behavior (Chen et al., 2018; Kiger et al., 2001; Leatherman and Dinardo, 2010; Tulina and Matunis, 2001). In contrast, female GSCs do not require this pathway, though it is active in the region of the adult female niche where it signals to cap cells and escort cells (Decotto and Spradling, 2005; López-Onieva et al., 2008). Interestingly, even though this signal is present in both adult male and female GSC niches, we find that this pathway is only activated in male GSCs and not female GSCs (Figures 1A and 1B). Activation of the JAK/STAT pathway often leads to an increase in the Stat92E protein, and this is commonly used as an assay for pathway activation (Chen et al., 2003; Yan et al., 1996). We observed a clear increase of Stat92E protein in male GSCs (Arrows in Figure 1A), as has been previously observed, but we failed to see an increase in female GSCs (Arrows in Figure 1B). In contrast, the somatic cells surrounding female GSCs do exhibit Stat92E expression, consistent with a role for this pathway in the cap cells and escort cells (Figure 1B).

To determine whether the differential response to the JAK/STAT pathway is due to the sex chromosome constitution of the germ cells, we examined Stat92E immunoreactivity in XX males. Interestingly, we found greatly reduced Stat92E staining in GSCs of XX males compared to controls (Figures 1C–1E, note that: that L3 larvae were analyzed since their morphology is more normal than XX *tra* adults). Previously, we had shown that JAK/STAT signaling is normally male-specific in the embryonic gonad and the pathway can be activated in XX males (Wawersik et al., 2005). To investigate whether embryonic XY and XX germ cells also showed a differential response to the JAK/STAT pathway, we quantified the Stat92E immunofluorescence signal in XX males. Indeed, embryonic germ cells in XX males exhibited a lower level of Stat92E immunoreactivity than did XY controls, although the difference was not as great as in larval gonads (Figure 1F). Lastly, as an independent assay of JAK/STAT response, we used the M5-4 enhancer trap, which is responsive to the JAK/STAT pathway (Gönczy and DiNardo, 1996). In wt testes, M5-4 is expressed in the hub and the hub-proximal germ cells (GSCs and early spermatogonia), and this is what we observed in XY control testes (Figure 1G). In contrast, we detected no M5-4 expression in the germline of XX male testes, although the expression in the hub remained similar to wt males, as expected (Compare Figures 1H to 1G).

For comparison, we also examined signaling through the Bone Morphogenetic Protein (BMP) pathway, which has been shown to be important for regulation of both the male and female GSCs (Schulz et al., 2004; Xie and Spradling, 1998). In contrast to what we found for the JAK/STAT pathway, we observed phosphorylated Mothers Against Dpp (pMAD), a downstream target of activated BMP signaling, in both control and XX male germ cells (Figures 1I–1J). Taken together, we conclude that the male germline niche is capable of signaling to both XY and XX germ cells, but that XX germ cells have a reduced ability to respond to the JAK/STAT pathway compared to XY germ cells.

### JAK/STAT activation rescues XX germ cells in a male somatic environment

We next wanted to determine whether decreased JAK/STAT signaling contributed to the germline defects observed in XX males. Inclusion of a dominant activated allele of the Drosophila JAK (*hopscotch*, *hop^Tum^*) (Hanratty and Dearolf, 1993; Harrison et al., 1995), in XX males significantly rescued germline depletion in 8.4% (n=83) of testes (Figures 2C, 2G). The presence of *hop^Tum^* also increased the JAK/STAT response (as judged by Stat92E immunoreactivity) in XX male germ cells (Compare Figures 2G to 2F). Rescued testes exhibited a large increase in the number of germ cells, and “DAPI-bright” germ cells were now restricted normally to the testis tip (Figure 2C), indicating that these were not tumors of early germline, but rather the XX germ cells of rescued testes were differentiating. Consistent with this, differentiating sperm were observed in most of the rescued testes (Figure S2). A similar result was obtained when we expressed an activated Stat92E (UAS-Stat92E^ΔNΔC^) (Ekas et al., 2010) specifically in the germline, which restored STAT expression (Figure 2D and 2H) and rescued the germline (Figure 2D, 13.15% of testes, n=38; Figure S2). The rescue of XX males by the JAK/STAT pathway occurred in an “all or none” manner; either XX males appeared fully rescued as shown, or similar to controls, without exhibiting intermediate phenotypes. Since restoring JAK/STAT signaling to the germline is sufficient to rescue at least a fraction of XX males, we conclude that decreased JAK/STAT signaling is an important contributing factor to the inability of XX germ cells to develop normally in a male somatic environment.

### Sxl activity represses the JAK-STAT pathway

Since a major difference between XX and XY germ cells is the expression of Sxl in XX germ cells, we next asked whether *Sxl* was responsible for the decreased JAK/STAT response in XX germ cells. Knocking down *Sxl* in the germline (*nos*>*Sxl^RNAi^*) was able to restore Stat92E immunoreactivity to GSCs in XX males (Compare Figures 3F to 3E), and rescue the germline defects to a similar extent (Figure 3C, 9.4% [n=96]) as was observed when the JAK/STAT pathway was activated directly. When we combined activation of the JAK/STAT pathway with *Sxl* knockdown in XX males (XX *tra*, *hop^Tum^*, *nos*>*Sxl^RNAi^*) no further increase in the percentage of germline rescue was observed (8.7% [n=45]; Figure S2). The absence of an additive effect leads us to conclude that activation of JAK/STAT and reduction of *Sxl* are acting in similar ways to rescue XX male germ cells, and that likely one major effect of reducing *Sxl* is to restore JAK/STAT activity to these germ cells.

**Figure 3:**
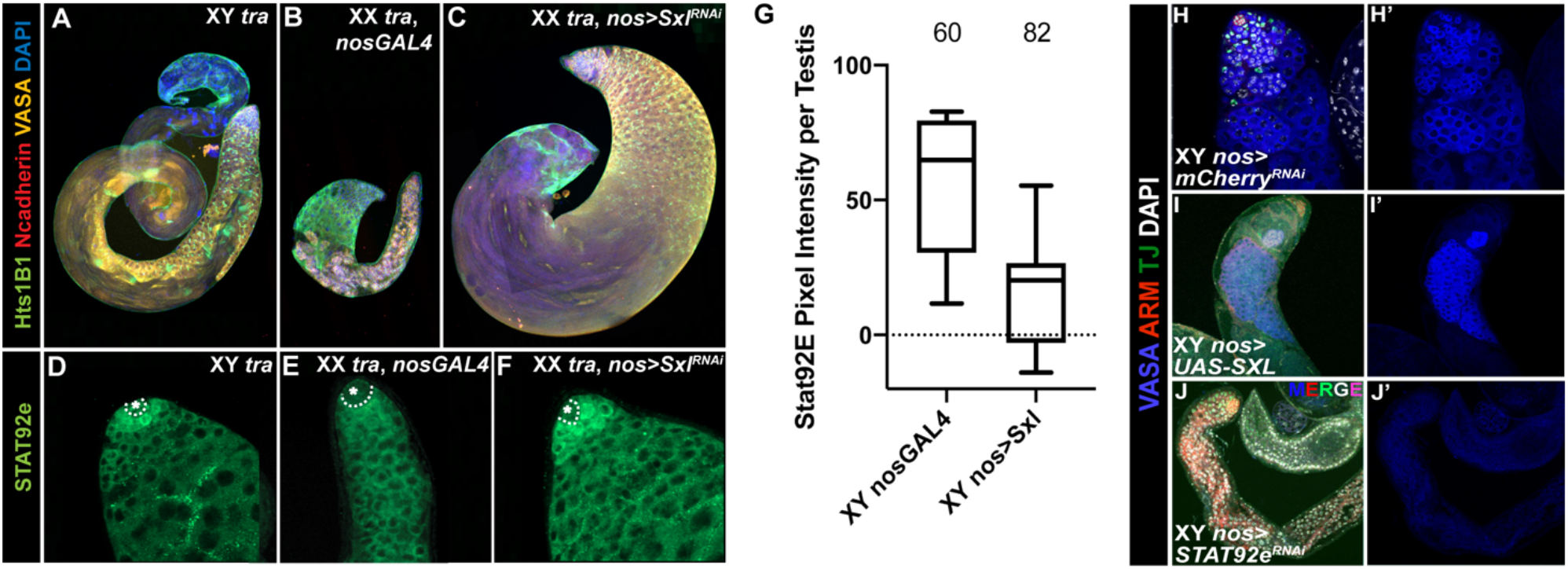
Sxl antagonizes the JAK-STAT pathway. (A-C) Hts1B1 (green), Ncadherin (red), Vasa (yellow), and DAPI (blue) in XY *tra* testes (A), XX *tra* >nos-Gal4 testes (B), XX *tra* testes expressing Sxl RNAi in germ cells, nos>Sxl^RNAi^ (C). (D-F) Stat92E (green) expression in XY *tra* testes (D), XX *tra* testes expressing nosGAL4 (E), and XX *tra* testes expressing Sxl RNAi in germ cells, nos>Sxl^RNAi^ (F). Hubs are indicated by dashed line in D-F. (G) Box and whiskers plot of the mean anti-Stat92E immunofluorescence pixel intensity per testis for XY testes expressing nosGAL4 (control) or XY testes expressing Sxl in germ cells, nos>Sxl. (H-J’) 2 week old adult testis immunostained as indicated. Germline overexpression of Sxl and depletion of Stat92E in the germline of testis cause similar germ cell loss.

We next asked if expression of *Sxl* was sufficient to reduce JAK/STAT signaling in otherwise normal XY testes. Quantification of fluorescence intensity indicated that Stat92E immunoreactivity was decreased in GSCs from testes expressing *Sxl* relative to wt testes (Figure 3G). We also observed a loss of GSCs and germline, similar to what is observed when JAK/STAT pathway activity is inhibited in the germline (Fig 3H–3J’). We conclude that Sxl expression is sufficient to reduce JAK/STAT signaling in otherwise wild type male GSCs.

### *Phf7* is a direct target of the JAK/STAT pathway in the germline

We next wanted to determine why sex-specific JAK/STAT signaling is important for male germ cells. Previously, we identified *Phf7* as being expressed male-specifically in the germline and being important for male fertility (Yang et al., 2012). *Phf7* is also toxic to female germ cells, and so it is also essential to repress it in the female germline. Interestingly, upregulation of *Phf7* was sufficient to rescue the germline defects in a fraction of XX males (Yang et al., 2012) similar to what we observed for activation of the JAK/STAT pathway or knockdown of *Sxl*. Therefore, we investigated whether *Phf7* was a direct target of the JAK/STAT pathway in the germline.

Previous RNA-Seq analysis by our lab and others indicated that *Phf7* utilizes alternative promoters in males and females, and that the upstream, male-specific promoter is repressed in a *Sxl*-dependent manner in females (Figure S3) (Primus et al., 2019; Shapiro-Kulnane et al., 2015). An interesting possibility is that the upstream promoter of *Phf7* might be regulated by the JAK/STAT pathway, which would explain both its preferential usage in males and the role of *Sxl* in regulating this promoter (Figure S3C). To test this idea, we first conducted qRT-PCR on wt testes, compared to testes where *Stat92E* was knocked down by RNAi. We observed that levels of the *Phf7* transcript from the upstream promoter were dramatically reduced in *Stat92E* germline knockdown testes compared to controls (Figure 4A). We also examined Phf7 protein expression using a hemagglutinin (HA)-epitope tagged *Phf7* genomic transgene that recapitulates male-specific Phf7 expression in both embryos and adults (Figures 4B, 4B’; Figure S4) and rescues the *Phf7* mutant phenotype (Yang et al., 2012) (Figure 5B). We observed a strong decrease in HA-Phf7 expression when *Stat92E* was depleted in the germline using two independent RNAi lines (Figure 4A and data not shown). Loss of HA-Phf7 expression was observed both in adult testes (Figures 4B–4C’) and male embryonic gonads (Figures 4D–4E’) upon *Stat92E* depletion. Together, the immunofluorescence and RT-PCR data indicate that the upstream promoter of *Phf7* is regulated by the JAK/STAT pathway and this has significant consequences on the production of the Phf7 protein.

**Figure 4.**
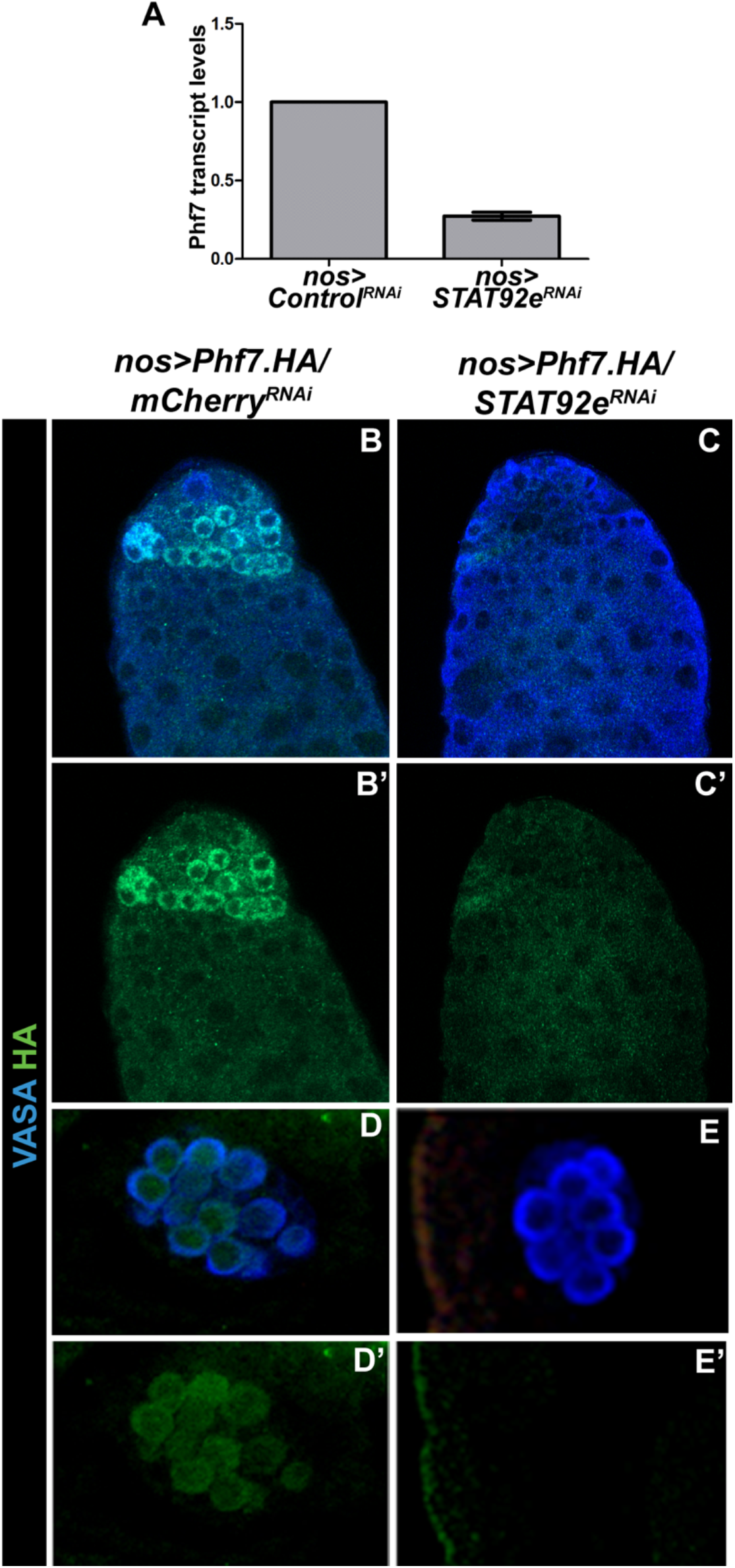
STAT regulates Phf7 expression in the germline. (A) qRT-PCR analysis of Phf7 transcript levels in control vs. germline Stat92E^RNAi^ adult testes. (B-D) Anti-HA immunolabeling to reveal expression from a HA-epitope tagged genomic *Phf7* transgene that rescues the *Phf7* mutant phenotype {Yang, 2012 #70}. (B, C’) Control and germline Stat92E^RNAi^ adult. (D, E’) Control and germline Stat92E^RNAi^ embryos. Note in all cases that knockdown of Stat92E leads to reduction in *Phf7* expression.

**Figure 5:**
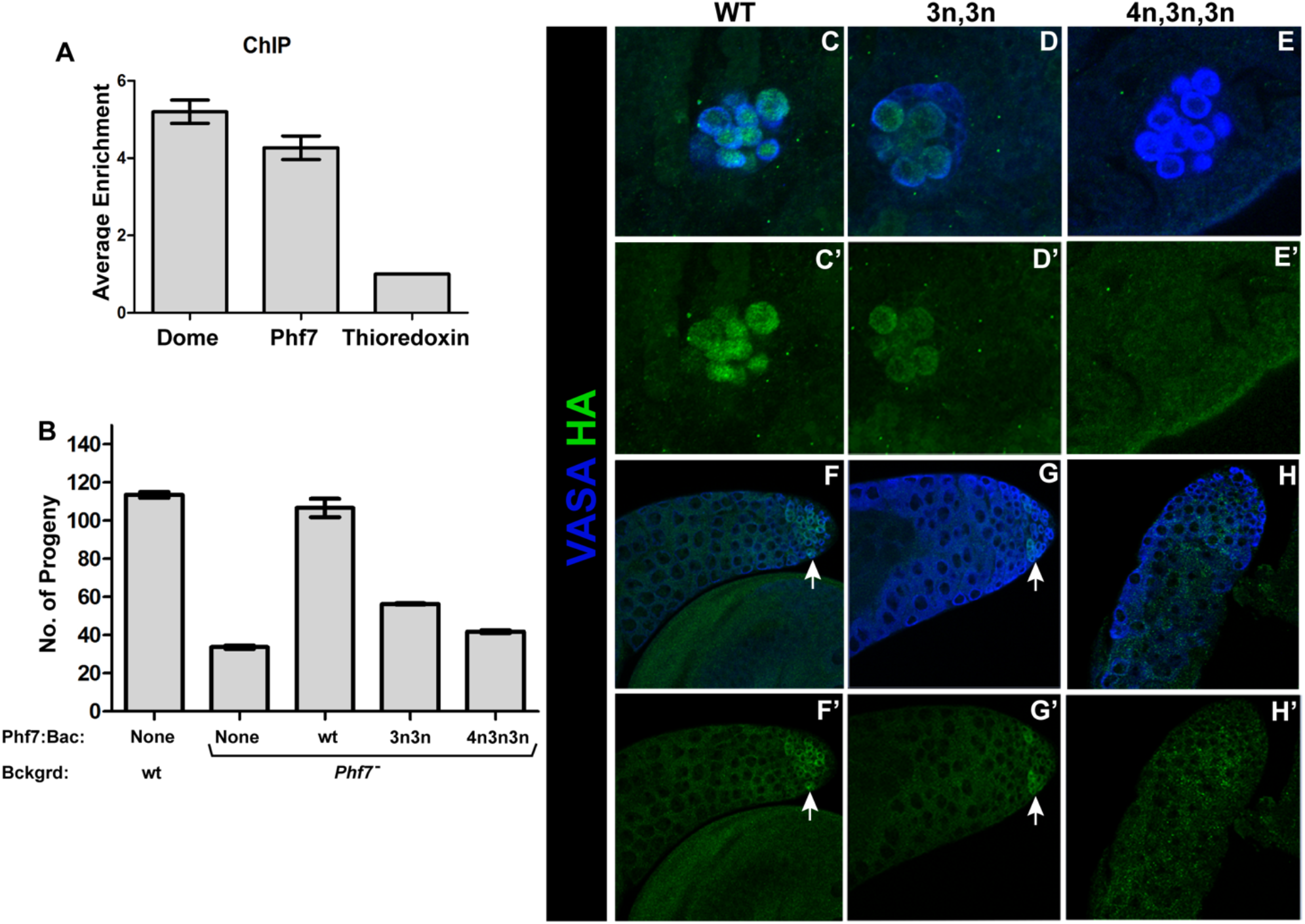
Stat92E is a direct transcriptional activator of *Phf7*. (A) Chromatin immunoprecipitation followed by PCR (ChIP-PCR) shows that STAT binds to Phf7 locus. Dome (Domeless) was used as positive control while *thioredoxin* was used as negative control. (B) Rescue of the *Phf7* mutant fecundity defect by *HA-Phf7* genomic transgenes. Genotypes are as indicated. 3n3n refers to the transgene with two Stat92E binding sites mutated and 4n3n3n refers to the transgene with all three Stat92E binding sites mutated. (C-H’) gonads immunostained to reveal expression of *HA-Phf7* genomic transgenes. (C-E) embryonic male gonads. (F-H) Adult testes. HA-*Phf7* transgenes are as follows: (C, F) wild type transgene, (D, G) transgene with two concensus Stat92E binding sites mutated, (E, H) transgene with all three concensus Stat92E binding sites mutated. Note that loss of two binding sites greatly reduces *Phf7* expression while loss of all three appears to eliminate *Phf7* expression.

To determine whether *Phf7* is a direct target of the JAK/STAT pathway, we first used a GFP-tagged *Stat92E* transgene (Venken et al., 2009) to conduct chromatin immunoprecipitation followed by PCR (ChIP-PCR). Anti-GFP ChIP-PCR revealed that DNA from the *Phf7* locus was enriched relative to a negative control (*thioredoxin*), and to a similar degree as a known Stat92E target (*domeless*) (Rivas et al., 2008) (Figure 5A). STAT proteins are known to bind to the consensus sequence TTCN_2-4_GAA (Rivas et al., 2008; Yan et al., 1996) with Drosophila Stat92E exhibiting a preference for a 3-nucleotide spacer (3N) (Yan et al., 1996). Interestingly, there are 3 consensus STAT sites downstream of the male-specific transcription start (exon 1 of the male transcript, Figure S3C), two with 3N spacing and one with 4N spacing. We mutated these sites within the context of the *HA-Phf7* transgene to determine their importance for Phf7 expression (Figure S3C). Mutation of the two 3N sites caused a dramatic decrease in HA-Phf7 immunostaining in both the embryonic and adult germ cells (Figures 5D, 5D’ and 5G, 5G’), while mutation of all three consensus STAT binding sites led to an absence of HA-Phf7 immunoreactivity (Figure 5E, 5E’ and 5H, 5H’). Finally, we determined the functional consequences of mutating the consensus STAT binding sites for *Phf7* function *in vivo*. The *HA-Phf7* genomic transgene is able to rescue the fertility defects observed in *Phf7* mutants (Yang et al., 2012) (Figure 5B). Mutation of either two or all three of the STAT sites in *Phf7* exon 1 resulted in a failure of the modified *HA-Phf7* transgene to rescue the decreased fertility of *Phf7* mutants. We conclude that *Phf7* is a direct JAK/STAT target in the male germline and that this regulation is essential for Phf7 expression and function.

### *Phf7* is repressed in female germ cells via its 5’UTR

The JAK/STAT pathway is an important regulator of the upstream, male-specific promoter of *Phf7*. However, our previous RNA-Seq analysis indicated that there is a significant level of *Phf7* expression from the downstream promoter present in females (Figure S3A), and this is also evident from consortium data (http://www.modmine.org/release-33/report.do?id=97001165). Given that we observe little expression of Phf7 protein in the ovary (Yang et al., 2012) (Figure 6), and that Phf7 expression is toxic to the female germline, we wondered if the *Phf7* transcript from the downstream promoter was subject to translational repression.

**Figure 6:**
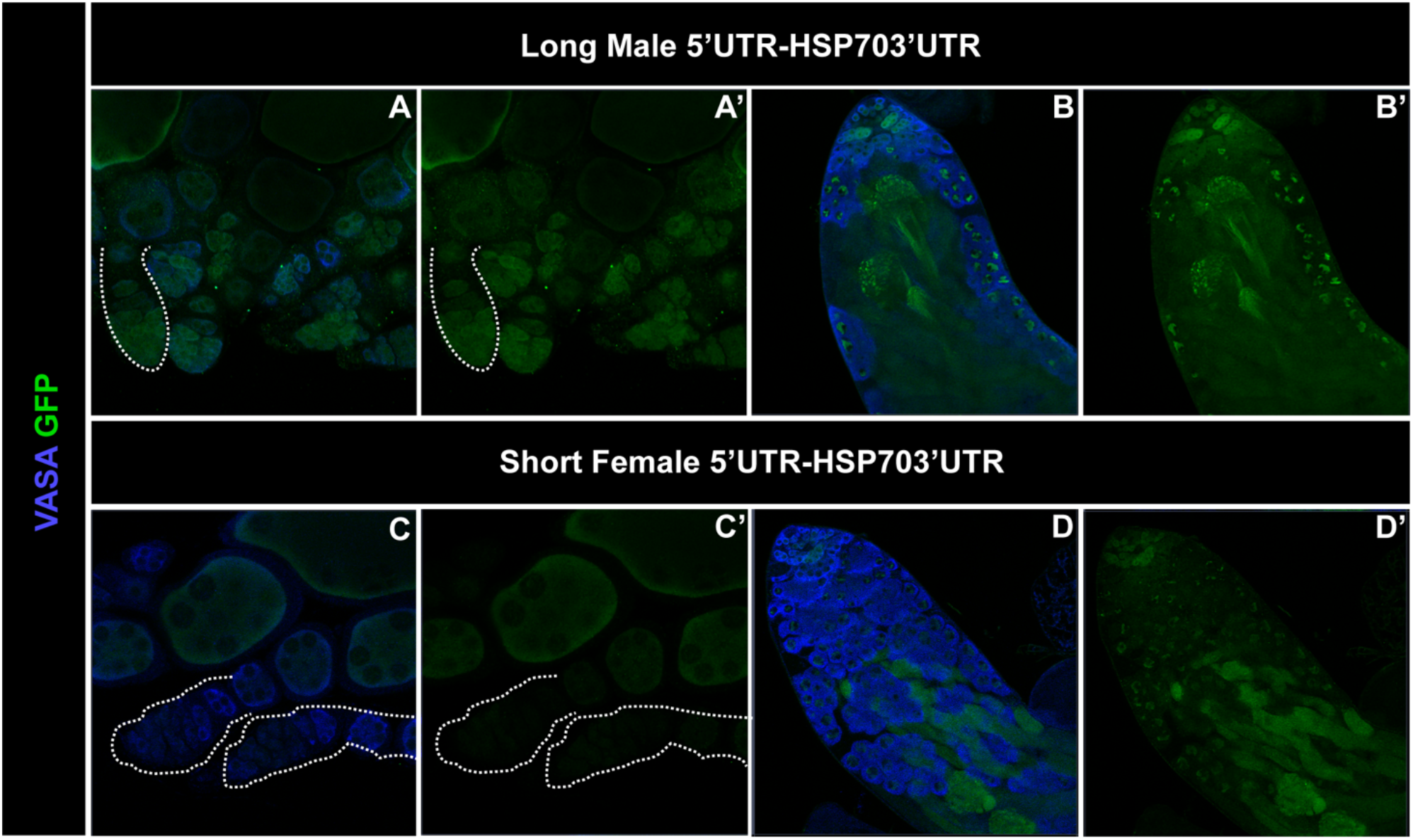
*Phf7* is repressed in female germ cells via its 5’UTR. Expression of UAS-GFP transgenes with either the longer, male-biased 5’UTR (A, A’ and B, B’) or the shorter, female-biased 5’UTR (C, C’ and D, D’) as indicated by anti-GFP immunostaining. Expression is shown in both ovaries (A, C) and testes (B, D). Note that expression of the short 5’UTR construct (C, C’) is much lower than expression of the long 5’UTR (A, A’) in the germaria of ovaries. There is also a difference in testes although some expression from the short 5’UTR (D, D’) persists in testes.

The *Phf7* mRNA from the male-biased, upstream promoter includes the entire female mRNA along with 170nt of additional 5’UTR. We constructed Gal4-responsive (UAS) GFP transgenes containing the 5’UTRs from either the long “male” transcript from the upstream promoter, or the shorter “female” transcript from the downstream promoter (Figure S3) and compared their expression when driven by *nos-Gal4* in the male and female germline. We found that, in the germarium of females, germline GFP expression was greatly reduced from the construct containing the female 5’UTR compared to the male 5’UTR (outlines, Figures 6A–6D’). Increased levels of GFP expression were observed in later egg chambers expressing the female 5’UTR, suggesting that repression of this message might be specific to the early germline. Interestingly, GFP expression was more similar between the male and female 5’UTR constructs in the male germline, with expression from the female 5’UTR construct still clearly visible. The female 5’UTR is a shorter version of the male 5’UTR but contains the same sequences and utilizes the same translation start sequence. Why the shorter, female 5’UTR should lead to repression in the ovary is unknown, but it is likely important to prevent expression of Phf7 protein in the undifferentiated female germline, where it is toxic (Yang et al., 2012).

### The relationship between *Phf7*, the JAK/STAT pathway and *Sxl*

Since the JAK/STAT pathway is upregulated upon loss of *Sxl*, which should lead to *Phf7* upregulation, we next wanted to determine whether increased Phf7 expression contributes to the defects observed in *Sxl* depleted XX germ cells. To determine if Phf7 is upregulated in XX germ cells lacking *Sxl* function, we examined expression of the *HA-Phf7* genomic transgene in animals where *Sxl* was knocked down in the germ cells (*nos*>*Sxl^RNAi^*). Indeed, we observed upregulation of HA-Phf7 immunoreactivity in the tumorous germline of *nos*>*Sxl^RNAi^* females, while HA-Phf7 was not observed in control females (Figures 7A–7B’). A similar result was observed examining *HA-Phf7* expression in *sans fille* (*snf*) mutant ovaries, which also lack Sxl (Shapiro-Kulnane et al., 2015). The germline “tumor” phenotype observed in *nos*>*Sxl^RNAi^* animals (Figure 7D) is similar to that observed in other germline-specific *Sxl* loss of function conditions (Schüpbach, 1985), but is different from the strong germline loss phenotype observed when *Phf7* is ectopically expressed in female germ cells (Yang et al., 2012). Thus, loss of *Sxl* in the female germline may lead to a lower-level expression of Phf7 than was the case when *Phf7* expression was ectopically driven in the germline.

**Figure 7:**
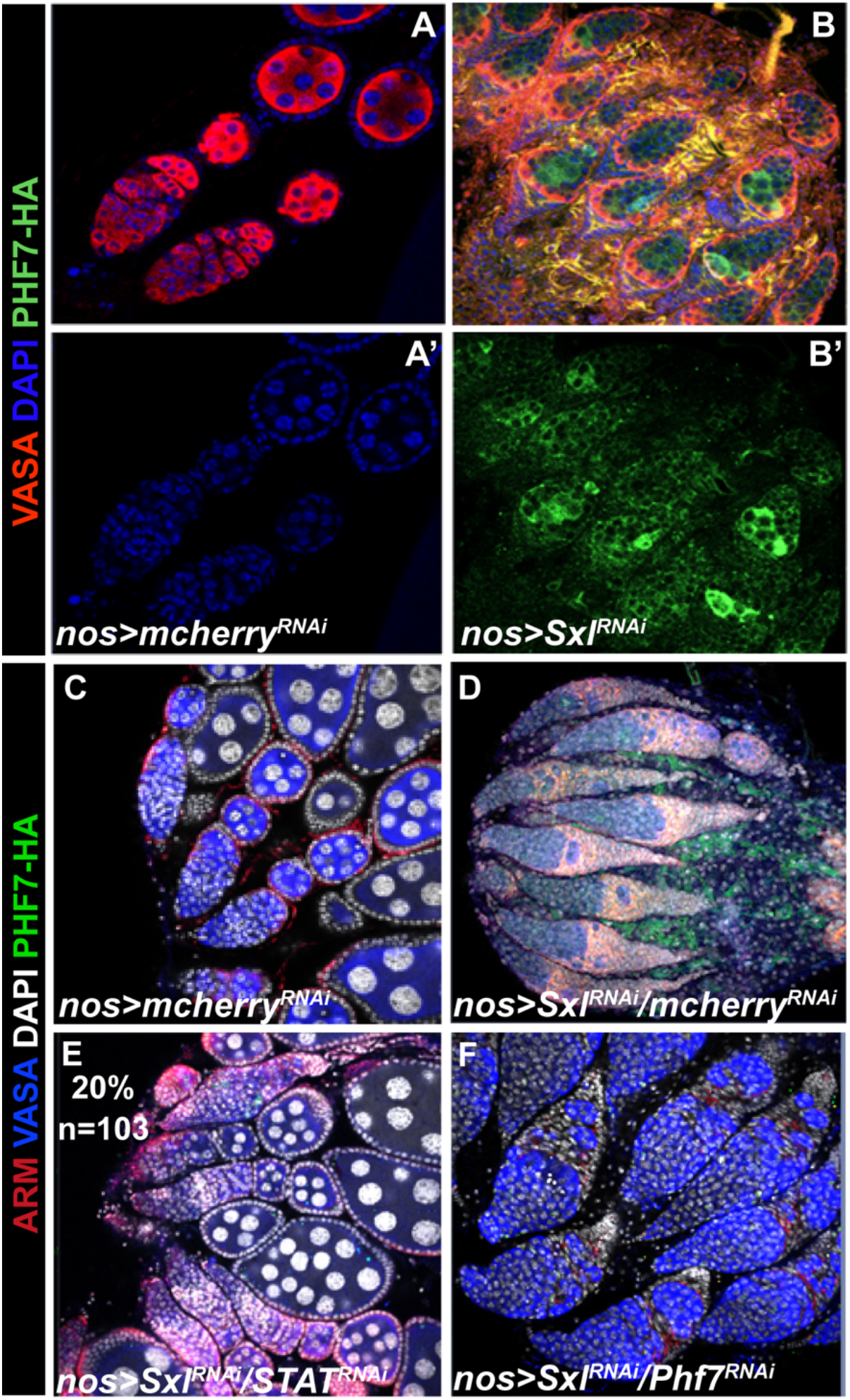
SXL promotes female sexual identity by repressing the JAK/STAT pathway. (A-B’) Germline depletion of Sxl results in tumorous ovary, which now exhibits Phf7 expression as judged by anti-HA immunostaining. (C-F) ability of different gene knockdowns to rescue the *Sxl* loss-of-function phenotype in the ovary. (C) control RNAi ovary. (D) Germline *Sxl* RNAi combined with control RNAi. Note the strong ovarian tumor phenotype. (E) *Sxl* RNAi combined with Stat92E RNAi. Note that these ovaries are rescued and appear similar to control ovaries (C). *Sxl* RNAi combined with *Phf7* RNAi. No rescue of the *Sxl* germline loss of function phenotype is observed.

We next wanted to determine if the increased Phf7 expression observed in *nos*>*Sxl^RNAi^* females was the main factor causing the *nos*>*Sxl^RNAi^* germline tumor phenotype. If this is the case, then blocking *Phf7* function should rescue the *nos*>*Sxl^RNAi^* mutant phenotype. However, we observed no such rescue and loss of both *Sxl* and *Phf7* appeared very similar or identical to loss of *Sxl* alone (Figure 7F). This is in contrast to what has been reported by the Salz lab using *snf* mutants which, again, lead to a loss of Sxl in the germline. They report that knocking down *Phf7* in the germline (*nos*>*Phf7^RNAi^*) was able to rescue the *snf* ovarian tumor phenotype and restore oogenesis (Shapiro-Kulnane et al., 2015). Due to the conflicting nature of these results, we repeated the experiment reported by Salz and colleagues using the same genetic reagents. However, in contrast to their observations, we saw no rescue of the *snf* ovarian tumor phenotype when *Phf7* was depleted in the germline. We then conducted similar experiments using a null allele of *Phf7* in combination with either the same *snf* mutants (snf^148^) or *nos*>*Sxl^RNAi^*. Again, we so no effect of loss of *Phf7* on the ovarian tumor phenotype caused by loss of *Sxl* or *snf* function (data not shown). We did, however, observe rescue of the *Sxl* loss of function phenotype when we knocked down *Stat92E* in the germline (Figure 7E). While *nos*>*Sxl^RNAi^* induces ovarian tumors in 100% of ovaries, simultaneous depletion of *Stat92E* was able to rescue egg chamber production in 20% of ovaries (N=103). This again indicates that regulation of the JAK/STAT pathway is a key function for Sxl in the female germline. However, while regulation of *Phf7* is an important aspect of why *Sxl* is essential in the female germline, it is clearly not the only factor leading to the defects caused by loss of *Sxl*.

## Discussion

Here we present data that provide new insights into germline sex determination and the regulation of male vs. female GSC identity. First, we find that one key function of the JAK/STAT pathway in GSCs is to promote male identity and directly activate expression of the male germline chromatin regulator Phf7. Further, we find that an important role for *Sxl* in female germ cells is to block the JAK/STAT pathway and prevent this signal from masculinizing the germline. Therefore, one key aspect of germline sexual identity is to regulate how GSCs respond to signals in their niche environment.

### The role of the JAK/STAT pathway in male GSCs

Different findings have led to different conclusions about the role of the JAK/STAT pathway in male GSCs. When STAT activity is removed from individual GSCs, they are lost rapidly from the niche, indicating a role in GSC identity or maintenance (Kiger et al., 2001; Tulina and Matunis, 2001). However, when STAT is removed from all GSCs, they exhibit defects in niche adhesion but can otherwise function as GSCs (Leatherman and Dinardo, 2010), although GSC loss is also observed (Tarayrah et al., 2015; Chen et al., 2018). The JAK/STAT pathway has also been implicated in aging of GSCs and their niche (Lenhart et al., 2019). One interpretation of these diverse data would be that the JAK/STAT pathway is important for specific aspects of male GSC function, such as regulation of cell adhesion and the cell cycle, but is not required for stem cell identity per se.

We propose a different role for the JAK/STAT pathway which is to regulate GSC sexual identity. Previously we reported that the JAK/STAT pathway is important for establishing male identity in the embryonic germline (Wawersik et al., 2005). Here we show that one defect observed in XX germ cells present in a male soma is that they exhibit reduced JAK/STAT signaling (Figures 2B, 2F). Further, activation of the JAK/STAT pathway can partially rescue these XX germ cells, promoting a male identity and progression into spermatogenesis (Figures 2C, 2G and 2D, 2H). Thus, we propose that the JAK/STAT pathway remains a key masculinizing signal for the germline throughout development and into adulthood. One possibility is that the JAK/STAT pathway regulates only GSC sex, and that other roles, such as regulating a specific set of cell adhesion proteins, represent downstream consequences of altering sexual identity. Alternatively, the JAK/STAT pathway could regulate GSC sexual identity and other aspects of GSC behavior independently.

One important way in which the JAK/STAT pathway promotes a male identity in the germline is by activating the male sex determination factor Phf7. Previously, we have shown that Phf7 is important for male identity in the germline and proper spermatogenesis (Yang et al., 2012). Phf7 likely promotes male germline identity by acting as an epigenetic “reader” and binding to histones methylated at position H3K4 (Yang et al., 2012; Yang et al., 2017; Wang et al., 2019). Phf7 is also toxic to female germ cells, making the sex-specific regulation of Phf7 extremely important. Here we show that the JAK/STAT pathway is a direct regulator of Phf7 expression in both embryos and adults. STAT protein can bind to the Phf7 locus (Figure 5A), and consensus STAT binding sites near the male-biased promoter are essential for proper male expression of Phf7 and its ability to function in spermatogenesis (Figures 5C–5H’; Figure 5B). Expression from the male-biased promoter is important in part because the transcript from the downstream, “female” promoter is subject to translational repression (Figures 6A–6D’). Thus, *Phf7* represents an important link between the JAK/STAT pathway and male identity in the germline.

### The role of Sxl in the germline

*Sxl* acts as a key regulator of sex determination in both the soma and the germline, and it is necessary and sufficient to confer female identity. However, the role of Sxl in the germline has remained mysterious. In the soma, Sxl regulates sexual identity through *tra* and dosage compensation through *msl-2* (Bell et al., 1988; Keyes et al., 1992; Bashaw and Baker, 1997; Kelley et al., 1997; Gebauer et al., 1998; Penalva and Sánchez, 2003), but these genes do not play a role in the germline. Instead, we have found that a key role of *Sxl* in the germline is to repress the JAK/STAT pathway in female germ cells.

Initially, only the male somatic gonad expresses ligands for the JAK/STAT pathway and is capable of promoting JAK/STAT activation in the germ cells (Wawersik et al., 2005). However, ligands for the JAK/STAT pathway eventually become active in the germarium of the ovary (López-Onieva et al., 2008), where they are important for the function or maintenance of the somatic escort cells (Decotto and Spradling, 2005). *Sxl* acts to repress JAK/STAT response in the female germ cells and thereby prevents activation of male-promoting factors such as Phf7. Somatic cells of the ovary such as the escort cells are still able to respond to these ligands and activate the JAK/STAT pathway, even though they also express Sxl. How Sxl is able to repress the JAK/STAT response in a germline-specific manner remains unknown, although the levels of Sxl appear higher in the GSCs than in the surrounding soma (Figure S1). Initially, our RNA-seq data revealed that the RNA for Drosophila JAK (*hop*) is differentially spliced in male vs. female adult gonads, and this differential splicing could possibly disrupt the Hop open reading frame (Figure S5). However, depletion of *Sxl* from the germline did not influence sex-specific splicing of *hop* (Figure S6) and extensive experimentation failed to reveal a role for *Sxl* in modifying germline splicing of *hop*. Thus, this is unlikely to be the mechanism for Sxl regulation of the JAK/STAT pathway in the germline. However, the fact that an activated Hop (hop^Tum^) can partially rescue the germline in XX males indicates that Sxl is repressing the pathway at the level of Hop or above. Interestingly, RNA for the JAK/STAT receptor *domeless* was identified in a pull-down experiment with Sxl, suggesting this could be a relevant target for regulation (Ota et al., 2017).

Our data support a model where the JAK/STAT pathway is important for activating male identity in the germline, and expression of male genes such as Phf7, while this pathway is repressed in female germ cells by Sxl. Loss of Sxl from the female germline leads to both upregulation of JAK/STAT signaling, and inappropriate expression of Phf7 (Figure 7) (Shapiro-Kulnane et al., 2015). In addition, suppression of the JAK/STAT pathway can partially rescue loss of *Sxl* in the female germline (Figure 7E). Thus, regulation of the JAK/STAT pathway is one key aspect of how *Sxl* promotes female germline identity. However, we observed no ability for loss of *Phf7* to rescue loss of *Sxl* from the female germline. This is in contrast to previously published results where loss of *Phf7* was shown to rescue the female germline in *sans fille* mutants, which also primarily affects the germline by disrupting Sxl expression (Shapiro-Kulnane et al., 2015). As discussed above, we have now reduced *Phf7* function by both RNAi and using null *Phf7* mutants, in both *Sxl* and *sans fille* loss of function backgrounds and observed no rescue or modification of the germline defects present. We conclude that, while regulation of *Phf7* by the JAK/STAT pathway and *Sxl* is clearly important for proper germline sexual development, ectopic expression of *Phf7* is not the only defect present in *Sxl* mutant female germ cells; there must be additional targets for regulation by *Sxl* and JAK/STAT that are disrupted in *Sxl* mutants. In support of this view, loss of Stat from the male germline has a more severe phenotype than loss of *Phf7* (Figures 3H–3J’) (Yang et al., 2012). Previously, we have shown that expression of another male-promoting factor in the germline, Tdrd5l, is regulated by Sxl (Primus et al., 2019). While this regulation appears to be, at least in part, via Sxl acting on the *Tdrd5l* mRNA to influence levels of Tdrd5l protein, it is possible that *Tdrd5l* is also be regulated at the transcriptional level as an additional target of the JAK/STAT pathway.

It is intriguing that *Sxl* acts as a negative regulator of the JAK/STAT pathway in both the soma and the germline but does so in different ways. In the soma, *Sxl* activates an alternative splicing cascade that leads to splicing of *dsx* in the female mode, creating the DSXF protein, while the DSXM protein is produced in males by default. An important sex-specific trait in the embryonic gonad is that male somatic cells produce ligands for the JAK/STAT pathway which activate JAK/STAT signaling specifically in male germ cells and this is regulated in a manner dependent on *dsx* (Wawersik et al., 2005). Thus, in addition to being a negative regulator of JAK/STAT signal reception in the germline, *Sxl* acts as a negative regulator of JAK/STAT ligand production in the soma. Together, these independent aspects of regulation by *Sxl* combine to ensure that the masculinizing effects of the JAK/STAT pathway are restricted to male germ cells.

### Germline sex determination

An important conclusion from this work is that germline sex determination regulates how GSCs communicate with their surrounding stem cell niche. Germline sex determination is regulated by both germline autonomous cues, based on the germline sex chromosome constitution, and non-autonomous signals from the soma. We have shown that the autonomous cues, acting through *Sxl*, regulate how signals from the niche are received and interpreted by the GSCs. In both the testis and ovary GSC cell niches, the JAK/STAT pathway is important for regulating somatic cells like the cyst stem cells in the testis (Leatherman and Dinardo, 2008; Sheng et al., 2009; Sinden et al., 2012) and the escort cells in the ovary (Decotto and Spradling, 2005). However, this pathway is only required in the male GSCs, and not female GSCs (Kiger et al., 2001; Tulina and Matunis, 2001; Decotto and Spradling, 2005). We propose that it is essential to block JAK/STAT signaling in female GSCs to prevent their exposure to this masculinizing signal. Indeed, activation of the JAK/STAT signal is sufficient to promote male identity in XX germ cells (Figures 2A–2H), and removal of STAT is sufficient to partially rescue the defects observed in XX germ cells that have lost *Sxl*. Thus, a key aspect of how *Sxl* promotes female identity in the germline is to prevent female GSCs from being masculinized by activators of the JAK/STAT pathway present in the niche environment.

It is important to note that, when we refer to the sex chromosome genotype affecting germline “sex determination”, this could result from any contribution of sex chromosome genotype to successful spermatogenesis or oogenesis. For example, if dosage compensation is incomplete or non-existent in the germline, then the presence of two X chromosomes will lead to increased X chromosome gene expression, which may be incompatible with male germline differentiation. Similarly, a single X chromosome dose may be incompatible with oogenesis. It is also possible that the number of X chromosomes present in the germline has additional affects besides the presence or absence of Sxl expression. While XX germ cells present in a male soma exhibit severe atrophy and loss (Figure 2B), the expression of Sxl in the male germline has a much weaker phenotype (Figures 3 I, 3I’). Thus, there may be additional consequences of sex chromosome genotype on germline function beyond that which is controlled by *Sxl*. A better understanding of what germline sexual identity means in Drosophila, in particular at the level of whole-genome gene expression levels, is required before we can assess the true contribution of germline sex chromosome constitution to germline sex determination. Further, how the effects of X chromosome number on germline sexual development in Drosophila relate to infertility observed in patients with Disorders of Sexual Development (DSDs) such as Klinefelter’s and Turner’s Syndromes remains to be investigated.

## STAR 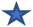 METHODS

Detailed methods are provided in the online version of this paper and include the following:

- KEYRESOURCESTABLE
- CONTACT FOR REAGENT AND RESOURCE SHARING
- Materials and Methods detail

- Fly Stocks
- Immunostaining
- Antibodies
- Quantifying Stat92E pixel intensity
- Fecundity test
- PHF7 UTR Construct
- Construct, mutagenesis and BAC recombineering
- STAT92e binding site recombineering
- galK positive/negative selection
- Electroporation [and Induction] of SW102
- Chromatin Immunoprecipitation (ChIP)

### KEY RESOURCES TABLE

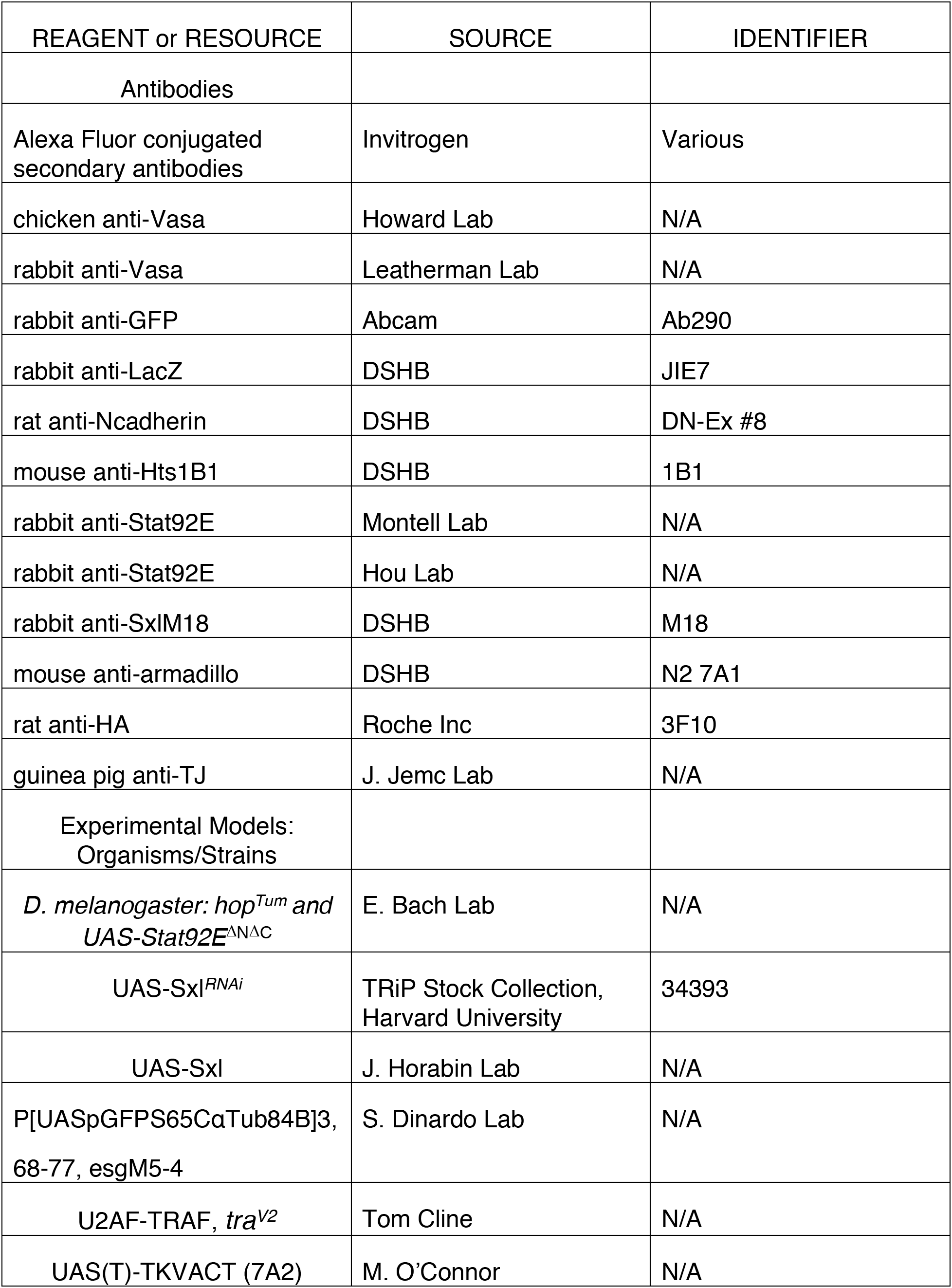
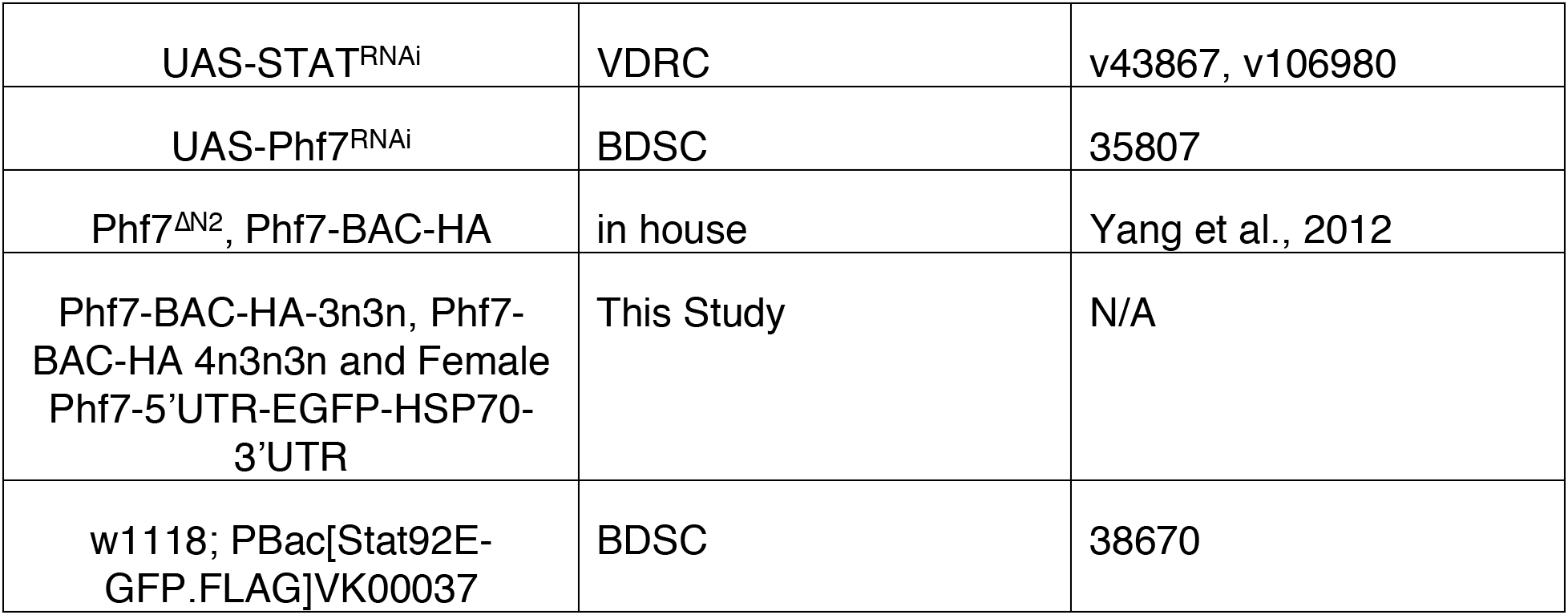

### CONTACT FOR REAGENT AND RESOURCE SHARING

Further information and requests for resources should be directed to and will be fulfilled by the Lead Contact, Mark Van Doren.

### Materials and Methods detail

#### Fly Stocks

The following fly stocks were used (unspecified stocks are from the Bloomington Stock Center): OregonR, *w^1118^* (as wild-type), *tra^1^*, Df(3L) st-j7 (*tra*^Δ^), *tra2^B^*, Df (trix) (*tra2*^Δ^, nanos-GAL4-VP16 (on Chromosome II or III). nanos-GAL4-VP16 driving UAS-Sxl were crossed and raised at 29C. Newly eclosed nos>Sxl flies were assayed for Stat92E levels while hub-expansion and GSC loss phenotypes were assayed in flies aged 7-10 days post eclosion.

#### Immunostaining

##### Embryo and larvae collection

Embryos and larvae were collected from agar plates, rinsed with PBT (PBS and 0.1% Triton) in a cell strainer. Embryos were dechorionated with 50% bleach for 5 minutes, rinsed with PBT and transferred to a vial, then fixed (8mL heptane, 0.25 mL 37% paraformaldehyde, and 1.75 ml PEMS [100mM PIPES, 2mM MgSO_4_, 1mm EGTA]) for 20 minutes with gentle agitation. The aqueous bottom layer was removed and 10mL of methanol was added to the vial that was then shaken for 30-60 seconds. Most of the liquid and material at the interface was removed and embryos were washed with methanol three times. Embryos were rehydrated with PBT with two, 5-minute washes and a final 30-minute wash before being immunostained.

Newly eclosed larvae were collected and fixed much like embryos with some changes. They were quickly rinsed with 50% bleach for 1 minute. After the initial 20-minute fixation they were transferred to 4.5% PFA (containing 0.1% Tween) for an additional 5 minutes. They were processed through a dehydration series in methanol/PBS and stored. For immunostaining they were rehydrated through a series of methanol/PBS solutions.

##### Embryo Immunostaining

Embryos were immunostained as follows: Samples were washed twice with PBT (for embryos Tween was used). Samples were briefly sonicated at lowest settings, resuspended in PBT and washed 3 times. Embryos were blocked with PBT containing 0.5% BSA and 2% normal goat serum for 30 minutes. Primary antibodies were added to the blocking solution for primary incubations lasting between 4 hours at room temperature to overnight at 4°C. Stat92E antibody solutions were incubated for 1.5 days, alternating room temperature and 4°C. Secondary antibody stains were performed following two PBT washes with antibodies diluted in the blocking solution above and incubations from 3 hours at room temperature to overnight at 4°C. Samples were washed twice with PBT and then positioned on a slide in PBS. The PBS was wicked away and samples were mounted in DAPBO.

Embryos and larvae were genotyped using transgenic balancer chromosomes and sex chromosome constitution was determined using paternal (P[Dfd-lacZ-HZ2.7]) or (P[Dfd-YFP]). Adult sex chromosome constitution was determined by the Y chromosome insertion Bs or by staining with msl1 antibodies that label the X chromosome of XY (but not XX) flies.

##### Immunolocalization on adult gonads

Day 1

1. Dissect tissue in 1X PBS. Transfer to a 1.5 ml tube containing 1X PBS. If it’s a quick dissection <20mins. You don’t need ice. If you will need more time, keep samples on ice.
2. Remove PBS and add fixative.
3. For Testes: Fix in 4.5% formaldehyde in PBTx, 20-30 min at room temp on nutator. 1 ml fixative: 125 ul formaldehyde 36% + 875 ul PBTx For Ovaries: Fix in 5.14% formaldehyde in PBTx, 10-15 min at room temp on nutator. 1 ml fixative: 100 ul formaldehyde 36% + 600 ul PBTx PBTx: PBS with 0.1% triton
4. Rinse twice in PBTx - Wash twice 10 min in PBTx. Not getting rid of fix will affect your immunostain.
5. Block at least 30 mins in PBTx + 0.5% BSA + 2% NGS (1 ml BBTx + 20 ul NGS). You can leave tissue in block overnight. Leaving the sample over the weekend (*adult gonads) hasn’t been shown to affect future steps.
6. Primary antibody overnight at 4°C (or 2h at room temp) in PBTx + 0.5% BSA + 2% NGS (0.5 ml BBTx + 10 ul NGS). For best images, do an overnight primary incubation. 300ul volumes of primary antibody are commonly used.

Day 2

7. Rinse twice in PBTx - Wash twice 10 min in PBTx
8. Secondary antibody 3h at room temp (or overnight at 4°C) in PBTx + 0.5% BSA + 2% NGS. (0.5 ml BBTx + 10 ul NGS)
9. Rinse twice in PBTx - Wash twice 10 min in PBTx Store in PBS at 4°C
10. Mount on slide in PBS. Dry PBS and replace by DABCO or Vectashield.

##### Immunolocalization on larval gonads

Day 1

1. Dissect tissue in 1X PBS. Transfer to a 1.5 ml tube containing 1X PBS. If it’s a quick dissection <20mins. You don’t need ice. If you will need more time, keep samples on ice.
2. Remove PBS and add fixative.
3. Fix in 5.14% formaldehyde in PBTx, 10-15 min at room temp on nutator. 1 ml fixative: 100 □l formaldehyde 36% + 600 ul PBTx PBTx: PBS with 0.1% triton
4. Rinse twice in PBTx - Wash twice 10 min in PBTx. Not getting rid of fix will affect your immunostain. The larval gonads will float. Be patient.
5. Block at least 30 mins in PBTx + 0.5% BSA + 2% NGS (1 ml BBTx + 20 ul NGS). You can leave tissue in block overnight. Leaving the sample over the weekend (*adult gonads) hasn’t been shown to affect future steps.
6. Primary antibody overnight at 4°C (or 2h at room temp) in PBTx + 0.5% BSA + 2% NGS (0.5 ml BBTx + 10 ul NGS). For best images, do an overnight primary incubation. 300ul volumes of primary antibody are commonly used.

Day 2

7. Rinse twice in PBTx - Wash twice 10 min in PBTx
8. Secondary antibody 3h at room temp (or overnight at 4°C) in PBTx + 0.5% BSA + 2% NGS. (0.5 ml BBTx + 10 ul NGS)
9. Rinse twice in PBTx - Wash twice 10 min in PBTx Store in PBS at 4°C
10. Mount on slide in PBS. Dry PBS and replace by DABCO or Vectashield. Stat92E antibody solutions were incubated for 1.5 days, alternating room temperature and 4°C.

#### Antibodies

Antibodies used (unspecified stocks are from the Developmental Studies Hybridoma Bank): chicken anti-Vasa (1:10000, Howard Lab), rabbit anti-Vasa(1:1000, Leatherman Lab) chicken anti-GFP(1:5000, Abcam), rabbit anti-LacZ (1:10000), rat anti-Ncadherin (1:12), mouse anti-Hts1B1 (1:4), rabbit anti-Stat92E (Montell Lab) (1:800, used in larvae and adults), rabbit anti-Stat92E (Hou Lab) (1:50, used on embryos), rabbit anti-SxlM18 (1:20), mouse anti-armadillo (1:100, DSHB), rat anti-HA (1:100, Roche Inc, used in embryos and adults), guinea pig anti-TJ 1:1,000 (J. Jemc).

Rescue of XX *tra* mutant testes were assessed by DAPI staining with more intense DAPI restricted to the germline in the apical tip of the testes. Visualization of sperm bundles was used to categorize testes as ‘differentiation to sperm’. Antibodies such as rat anti-Ncad or mouse anti-Hts1B1 allowed for this visualization.

#### Quantifying Stat92E pixel intensity

Confocal slices were analyzed with ImageJ software to attain average pixel intensity per GSC (multiple confocal slices). Background staining in each testis was measured and subtracted from the GSC average pixel intensity. The GSCs of a testis were then averaged together, creating a data point. Data points were also averaged to create a testis average for each genotype.

#### Fecundity test

For males, newly eclosed single test males were mated to 15 virgin Oregon-R females for 1 week and number of progenies produced was recorded 14 days after cross started.

#### Phf7 UTR Construct

5’, 3’ UTR of Phf7 and 3’UTR of HSP70 were cloned into pUASP vector using the listed primers. 5’UTR were cloned in front of EGFP. pUASpB is a modified version of pUASP (Rørth, 1998) including an attB site for phiC31-mediated integration. Phf7 UTR constructs were generated by cloning a flexible linker sequence followed by the ORF of *Phf7* (lacking a start codon) using the listed primers into UAS-*sfGFP* that had been linearized with Xba1 for 5’UTR cloning and with SpeI and Pst1 for 3’UTR cloning. Constructs were assembled using HiFi DNA Assembly (New England Biolabs).

EGFPEX0_fwd CCGCGGCCGCTCTAGCAGAAGCTCACAGGT
EX05UTR_rev CAAAAAGCGTTGAATTCCCGA
Male5UTRegfp_fwd-cgggaattcaacgctttttgAACAGCAGTTTTAGGCAATTCGTGCCAACG
Male5UTRegfp_rev-GCCCTTGCTCACCATG GATGCGTTTCCG
EGFPFemale5UTR_fwd CCGCGGCCGCTCTAGGATTTGTT
EGFPFemale5UTR_rev GCCCTTGCTCACCATG GATGCGTTTC
HSP70B3UTRSPE1Fw-GGG CCC ACTAGT ccaaatagaaaattattcagttcctg
HSP70B3UTR PST1Rv-AAT CG CTGCAG tatattctatttattaaccaagtagc

Constructs flies were injected into embryos and integrated via PhiC31 integrase-mediated transgenesis (done at BestGene Inc., California) into the same genomic location at P[CaryP]attP40 on Chromosome II. For germline-specific overexpression, male flies carrying UAS transgenes were mated with *nanos*-GAL4:VP16 virgin females (Van Doren et al., 1998) and the crosses were maintained at 29°C. The progeny was reared at 29°C until 3-5 days post-eclosion (unless otherwise specified).

#### Construct, mutagenesis and BAC recombineering

All Phf7 reporter construct was made in Phf7-BAC (CH322-177L19, BACPAC Resources Center, (Venken et al., 2009). It was recombineered to add 3xHA at C-terminus. The C-terminus tagged Phf7-BAC was inserted into PBac[yellow[+]-attP-3B]VK00033 and it successfully reports Phf7 expression (Yang et al., 2012).

The region of Phf7 carrying STAT92e binding sites was PCR amplified and cloned into PCR2.1-TOPO vector (Thermo Fisher). Site directed mutagenesis was carried out to mutate the 3 STAT92e binding sites. Galk recobineering method was used to clone back the fragment into Phf7-BAC-HA as described in (Warming et al., 2005).

For transgenesis these constructs were inserted into PBac[yellow[+]-attP-3B]VK00033 same as the control BAC for ideal comparisons of Phf7 expression.

#### STAT92e binding site recombineering

Region of Phf7 containing 3 STAT92e binding sites was PCR amplified and cloned into TA vector using primers below. Primers bind just outside where GALk primers end in Phf7 locus.

Phf7 TA 1 Fw-ggaatattattaagcaaatgatcaag
Phf7 TA 1 Rv-tgtgcaatgaattgtaacacaaa

Sites were mutated with primers below.

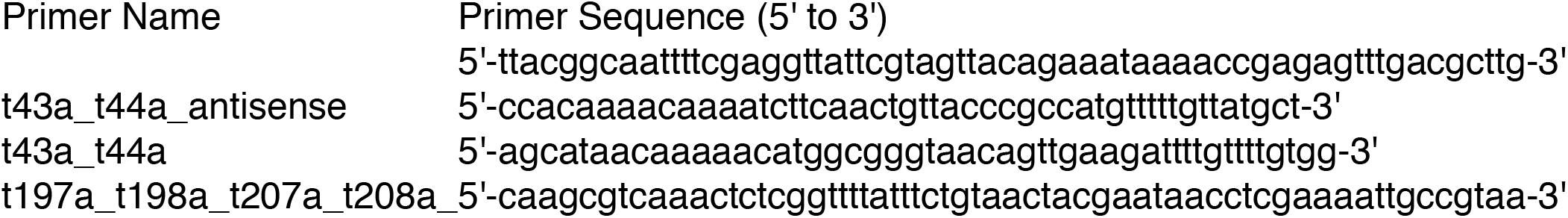

Galk sequence was amplified by primers below.

Phf7 STAT 3n4n GALk 1 Fw
tataacatatatatatatatagtttgtaaggtttatggggtcggaaacgcCCTGTTGACAATTAATCATCGGCA
Phf7 STAT 3n4n GALk Rv
tatatctacttgaaaaggcattaattagaacccctatacacatactttga
TCAGCACTGTCCTGCTCCTT

Mutated TA clone was used to substitute GALK in the vector. Protocol as outlined was used for galK recomineering (Warming et al., 2005)

#### Chromatin Immunoprecipitation (ChIP)

Chromatin Immunoprecipitation reaction was carried out in adult (3-5 day old) testis. About 200 pairs of testes (w1118; PBac[Stat92E-GFP.FLAG]VK00037) was subjected to ChIP according to (Tran et al., 2012). Real time PCR was performed using SYBR green mix on AB7300 system.

#### Chip primers

Primers were designed from this region:

gatttatatacattcggctagcataacaaaaacatggcgggtttcagttgaagattttgttttgtggctgtctcaaaaaacacgtgtgccaattcaatgtaaagtcctgcacaaataattaaacagttggaaaaacgcttaggggcaaccctactgccagagttg gcaagcgtcaaactctcggttttatttctgtttctacgaatttcctcgaaaattgccgtaacctggggcaaatgcgaattgaatgaaagttgcagtcctttattt

This covers all 3 STAT92e binding sites.

STAT-Phf7 chip Fw-ttcggctagcataacaaaaaca
STAT-Phf7 chip Rv-ggactgcaactttcattcaattc
Thiredoxin Chip Fw-caaattcgcatgctgtcagt
Thiredoxin Chip Rv-ggctgctggctgttctttac
domechipFw2-atttccattcaaggggttcc
domechipRv2-actggcgtgcatgtgtgta

---

**Supplemental Figure 1.**
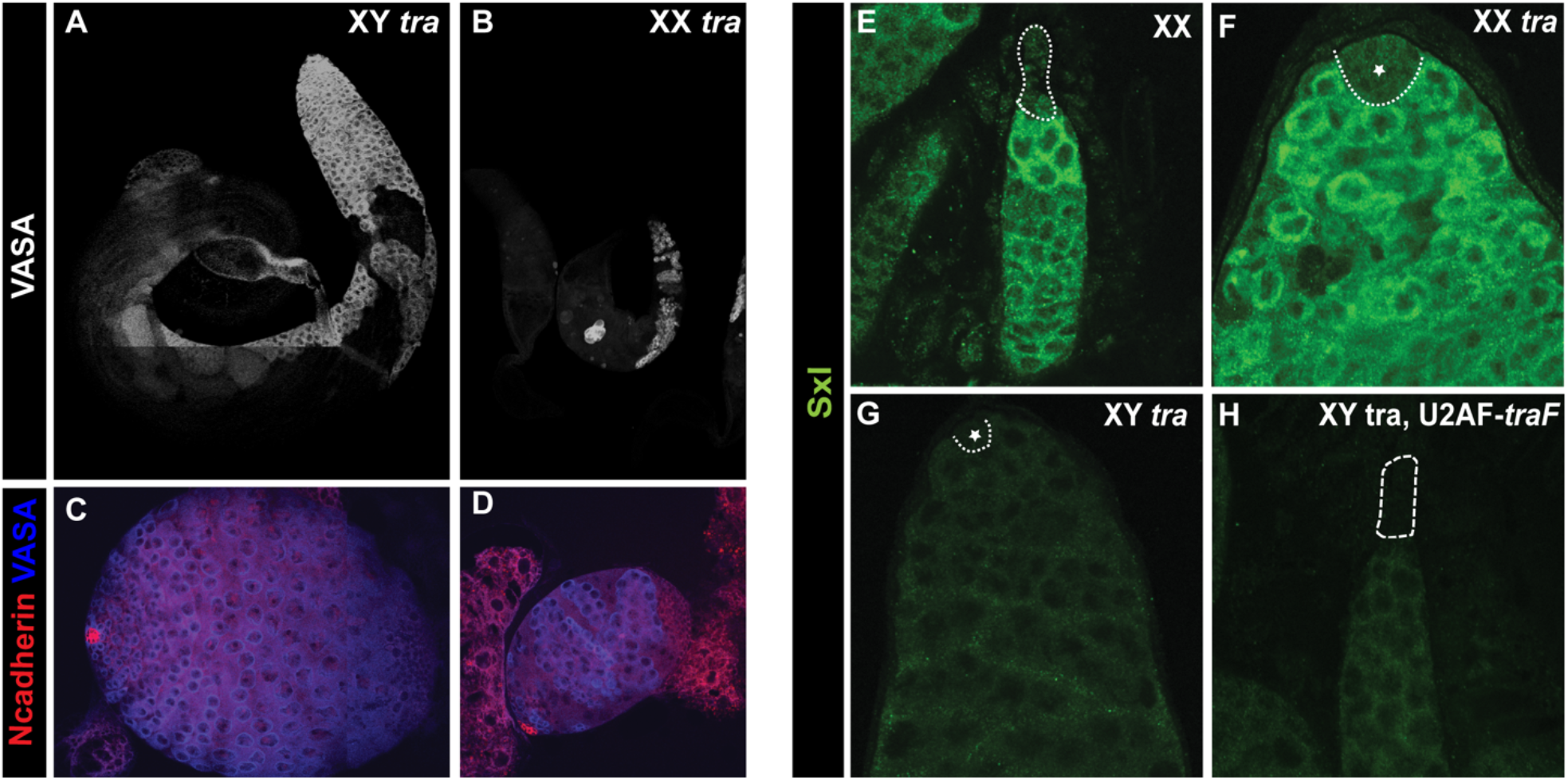
XX *tra* testes have depleted germline and show elevated Sxl levels compared to WT testes. (A, B) Adult XY *tra* (control) or XX *tra* (XX male) testes labeled with anti-Vasa to mark the germline. (C, D) L3 stage testes with germline labeled by Vasa (blue) and hubs labeled by Ncadherin (red). Note the greatly depleted germline in B and D. (E-H) Anti-Sxl staining of adult gonads. (E) Control XX ovariole, (F) XX *tra* testes, (G) XY *tra* (control) testes, and (H) XY tra, U2AF-traF (XY female) ovariole. Niches indicated by dashed line. Note that Sxl is highly expressed in XX germ cells and low in XY germ cells regardless of whether they are in a female or male somatic environment. * indicate hub.

**Supplemental Figure 2:**
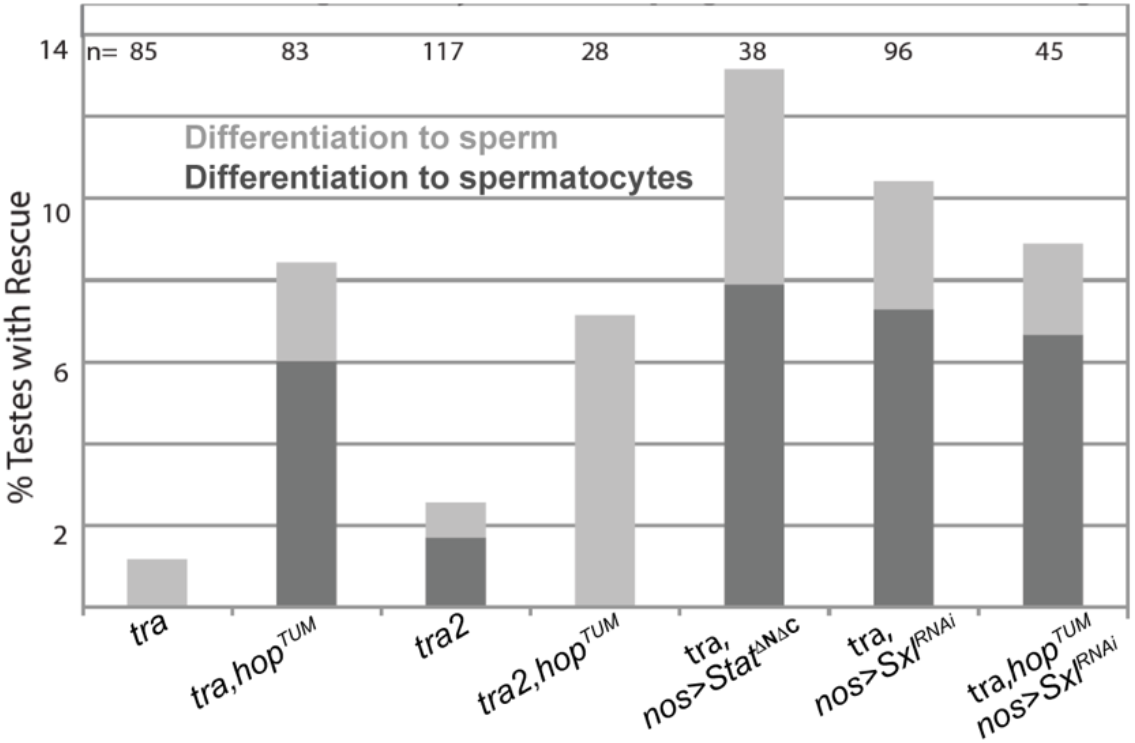
Quantification of the germline rescue of XX *tra* or *tra2* testes by activation of JAK/STAT or downregulation of Sxl. XX *tra* and *tra2* mutant testes exhibit a high % of atrophic testes with a lack of spermatocytes and mature sperm that can be rescued to different extents by activated JAK (*hop^Tum^*), expressing activated Stat92E in germ cells or knocking down Sxl in germ cells.

**Supplemental Figure 3:**
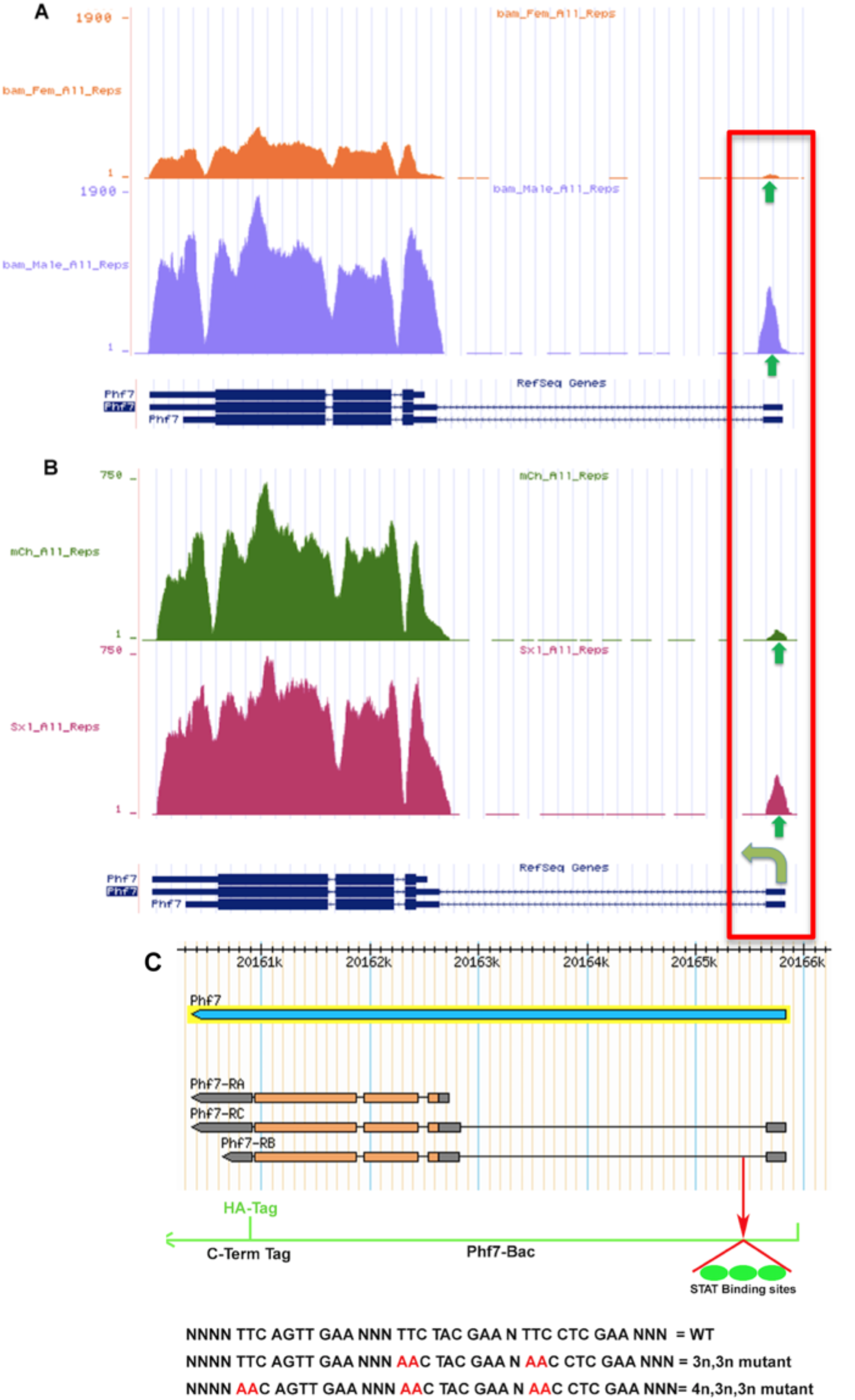
The Phf7 upstream promoter is male-biased. (A, B) RNAseq data from (Primus et al., 2019) showing reads mapped to the *Phf7* locus. *Bab of Marbles (Bam)* mutants were used to enrich for undifferentiated germ cells. (A) shows testes vs. ovaries. Note the strong increase in overall Phf7 expression in males, including from the upstream promoter (green arrows). (B) shows *Sxl* wild-type vs. knockdown. Note again the increase in expression from the upstream promoter in *Sxl* knockdown. (C) Position and sequence of 3 predicted STAT92e binding sites in intron 1 of *Phf7*. Sites were mutated by changing TTCN_2-4_GAA to AACN_2-4_GAA.

**Supplemental Figure 4:**
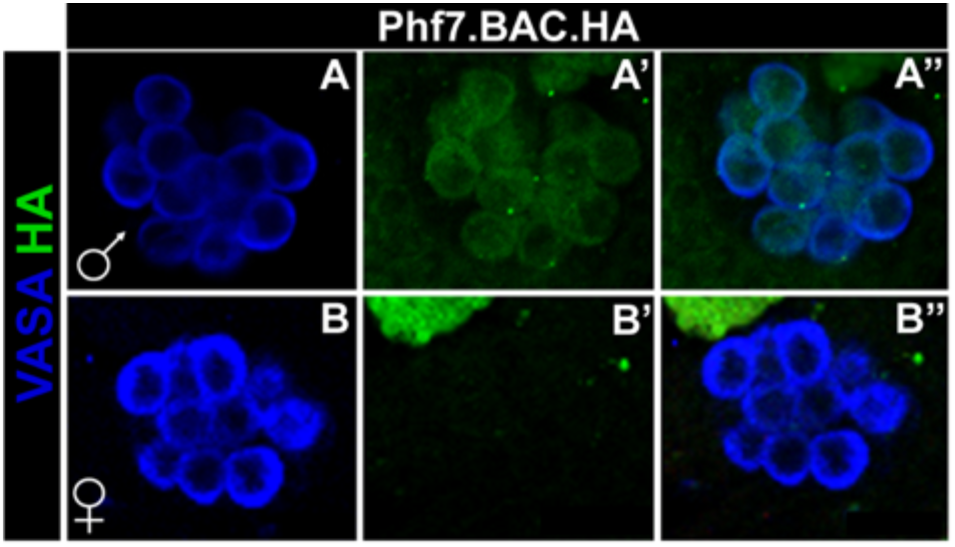
(A-B) Embryonic male (A) and female (B) gonads immunostained to reveal expression of the HA-tagged Phf7 genomic transgene. Note that *Phf7* is male-specific in embryonic germ cells as it is in adults.

**Supplementary Figure 5:**
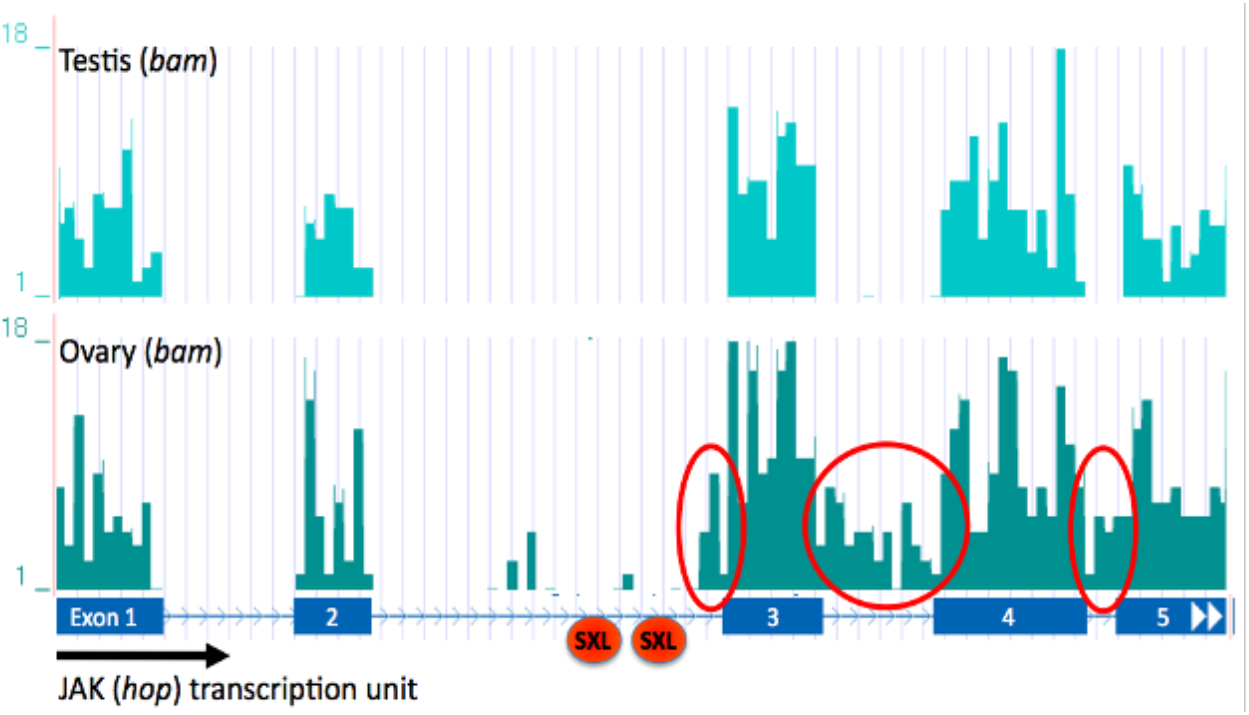
JAK (*hop*) is differentially spliced in male vs. female adult gonads. SXL is an RNA-binding protein that facilitates sex-specific alternative splicing of many targets, we analyzed RNA Sequencing data (http://www.genome.ucsc.edu, April 2006) of JAK/STAT pathway components to find candidate molecules. In female bam mutants containing only early germ cells, HOP appeared to have spliceforms that were not present in male bam mutants. Additionally, retention of HOP’s third intron would incorporate a nonsense codon into the transcript. This intron appears to be retained in transcripts of bam females. A nonsense codon could lead to nonsense-mediated decay and reduction of HOP transcript levels, thereby downregulating the JAK/STAT pathway.

**Supplementary Figure 6:**
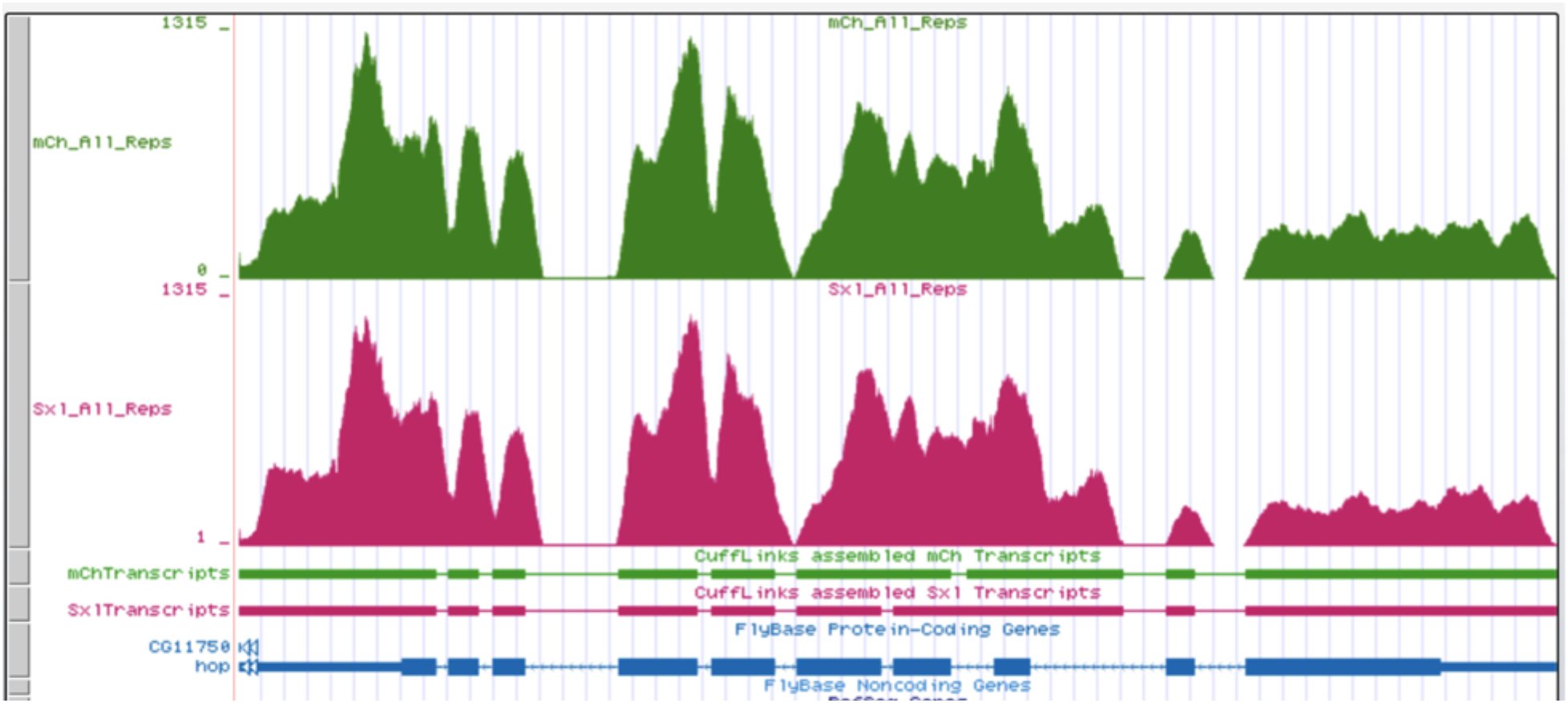
Depletion of *Sxl* from the germline did not influence sexspecific splicing of *hop*. RNAseq data suggest no changes in splicing when we down regulate Sxl in the germline.

## Notes

### Competing Interest Statement

The authors have declared no competing interest.

